# BMP signalling is critical for maintaining homeostasis of hair follicles and intestine in adult mice

**DOI:** 10.1101/174680

**Authors:** Aditi Nag, Pallavi Nigam, Abhishek L. Narayanan, Megha Kumar, Ritika Ghosal, Jonaki Sen, Amitabha Bandyopadhyay

## Abstract

BMP signalling play critical roles during embryonic development, however, its roles have not been extensively studied in the maintenance of adult tissue homeostasis. In this study we have used temporal knock out of *Bmp2* and *Bmp4* to uncover the role of BMP signalling in adult mice. We observed rapid changes in the adult hair follicles and the intestine within two days of depleting BMP signalling, which demonstrates the critical and acute requirement of BMP signalling in maintaining homeostasis in these tissues. In addition, our study demonstrates that while BMP signalling is required for maintenance of quiescence in telogen hair follicles, it is needed for survival and proliferation in late anagen hair follicles. Our study also reveals differential requirement of BMP signalling in differentiation of distinct intestinal cell types. In addition, Loss of BMP signalling rapidly promotes early neoplastic transformations in mouse intestine.

## 1. Introduction

Extensive studies on BMP signalling in the context of embryonic development have established that this signalling pathway plays pleiotropic roles during morphogenesis by regulating a variety of processes such as patterning, differentiation, proliferation and apoptosis. In adult animals, despite a steady state turnover of differentiated cells, homeostasis is maintained by the replacement of these cells by newly proliferated progenitor cells undergoing appropriate differentiation. It is thus, very likely that the same signalling pathways that govern embryonic aspects of morphogenesis will continue to regulate tissue homeostasis in adults. Studies using cell-type specific manipulation of BMP signalling in adult animals suggest that it is critical for maintaining tissue homeostasis in the neurogenic niches in the brain (Bond et al., 2014; Mira et al., 2010). However, no attempts have been made, thus far, to comprehensively assess the role of BMP signalling in maintaining various aspects of adult physiology. This would have to be done by manipulating this signalling pathway exclusively in adult animals, while allowing normal embryonic development to take place.

The hair follicle (HF) is considered to be a mini organ which develops through reciprocal interaction between the epithelial and the mesenchymal cells. BMP signals emanating from the mesenchymal cells influences the patterning and differentiation of the epithelial derived components of the HF (Botchkarev et al., 1999; Botchkarev and Paus, 2003; Rendl et al., 2005). Hair-follicle specific loss of function of BMP signalling, through mis-expression of Noggin driven by *KRT14* or *Msx2* enhancers led to the arrest of HF development at a very early stage (Kulessa et al., 2000; Plikus et al., 2008). This not only precluded the investigation of the role of BMP signalling in the adult HF but also failed to provide any insight into the role of this signalling pathway during later stages of HF development and cycling. On the other hand, cell type specific knock out of BMP receptor 1a (BmprIa) only produced cell-autonomous effects and thus did not provide sufficient insight about the different aspects of HF development regulated by BMP signalling (Yuhki et al., 2004). It was shown eventually that BMPs function as a “chalone” that inhibits hair growth during the resting phase of the hair cycle (Plikus et al., 2008). Recently, Genander *et al,* through the use of a variety of mouse molecular genetic tools demonstrated that BMP signalling modulates the specification of closely related HF progenitor cells (Genander et al., 2014).

Insight into the role of BMP signalling in intestine development has been primarily obtained through studies of Noggin misexpression driven by *Villin* or *Fabp1* enhancers and the intestinal epithelium specific knock-out of *Bmpr1a*. Although in these animals the manipulation of BMP signalling was initiated at embryonic stages, the changes in morphology, cell proliferation and crypt organization were observed much later in mice that were several weeks old (Batts et al., 2006; Haramis et al., 2004). Thus, it could not be ruled out that the observed changes in the intestine were due to chronic effects of loss of BMP signalling at embryonic stages which were manifested in the adult. Further, conditional knock-out of *Bmpr1a* driven by an absorptive-cell specific *Cre* (*Villin1::Cre*) did not produce any differentiation defect in the absorptive cell type nor in any of the secretory cell types other than the Paneth cells. Even in the Paneth cells only terminal differentiation was mildly affected (Auclair et al., 2007). While these studies clearly indicated that BMP signalling is necessary to supress ectopic crypt formation and polyp growth, the role(s) of BMP signalling are yet to be clearly elucidated in the context of intestinal homeostasis.

In the absence of any prior information on the possible contexts in which BMP signalling is likely to play a role in adult mice, we decided to inactivate BMP signalling ubiquitously in a temporally regulated manner. For this purpose we have used a mouse strain reported earlier, namely *Bmp2*^*C/C*^; *Bmp4*^*C/C*^; *R26CreER/+* (Yadav et al., 2012). These animals were injected with tamoxifen at six weeks of age to ubiquitously knock-out *Bmp2* and *Bmp4* exclusively in the adult stage. This strategy allowed us to assess the role of this signalling pathway specifically in adult animals avoiding any possible complications in interpreting the data arising out of the effects of depleting of BMP signalling at embryonic stages. The results obtained from our study suggest that BMP signalling is critically required for many aspects of tissue homeostasis in adult animals including HFs, intestine, adipose tissue and epidermis. In this report, we have extensively characterized the role of BMP signalling in regulating the homeostasis of HFs and the intestine. Our data suggests that BMP signalling plays distinct roles in proliferation, differentiation and cell survival in these tissues in adult mice.

## 2. Results

### 2.1. Depletion of BMP signalling in the adult mouse causes gross physiological derangements in the skin and intestine

To investigate the role of BMP signalling in adult mice, *Bmp2* and *Bmp4* were knocked-out in six week old *Bmp2*^*C/C*^; *Bmp4*^*C/C*^; *R26CreER/+* animals (Yadav et al., 2012) by intra-peritoneal injection of tamoxifen. We observed hair loss after the start of tamoxifen injection leading to formation of distinct bald patches within three weeks (compare Figures S1B to S1A). The animals were completely bald three months later (Figure S1C), showed significant loss of subcutaneous fat (Figure S1E) and occasionally developed sebaceous cysts (Figure S1F and G) and keratin cysts (Figure S1E, arrowheads). We also observed rectal prolapse in many of these animals (Figure S1H). We did not observe any of these phenotypes in age-matched control (*Bmp2*^*C/C*^; *Bmp4*^*C/C*^) animals injected with an identical regimen of tamoxifen. Henceforth the *Bmp2*^*C/C*^; *Bmp4*^*C/C*^; *R26CreER/+* animals will be referred to as “test animals” while the *Bmp2*^*C/C*^; *Bmp4*^*C/C*^ animals will be referred to as “control animals”. Based on the observed gross defects as well as existing literature about the involvement of BMP signalling pathway in morphogenesis of HFs and the small intestine (Auclair et al., 2007; Batts et al., 2006; Haramis et al., 2004; Plikus et al., 2008; Schneider et al., 2009; Yuhki et al., 2004) we decided to investigate the role of BMP signalling in depth in the context of the adult HF and small intestine, specifically jejunum. Although all of the gross defects mentioned above were observed in both female and male animals, we have carried out all the molecular and histological analyses of these phenotypes in male animals to avoid the confounding effects of hormonal changes in females that may additionally influence hair cycling (Foitzik et al., 2003; Oh and Smart, 1996).

### 2.2. Effect on proliferation and cell death upon depletion of BMP signalling in hair follicle and intestine of adult mice

We injected control and test male mice with tamoxifen and harvested skin and intestine samples every alternate day beginning with the third day up to the ninth day after the first injection. Domains of expression of *Bmp2*, *Bmp4*, Bmp receptors and the domains of BMP signalling in the HFs and the intestine are well documented (Haramis et al., 2004; Lee and Tumbar, 2012; Torihashi et al., 2009). To confirm that administration of tamoxifen was effective in the HFs and intestine, we examined the presence of pSMAD 1/5/8 immunoreactivity in these tissues. In both the tissues the reduction in pSMAD 1/5/8 immunoreactivity is evident after three days of tamoxifen injection (Figure S1I – S1L) while it was undetectable in both the tissues by the fifth day (Figure 1A-D). Studies have shown that reduction of BMP signalling causes induction of anagen and *de novo* proliferation in resting HFs (Genander et al., 2014; Plikus et al., 2008) while we observed loss of hair upon depletion of BMP signalling. To reconcile our observation with published reports, we investigated the status of cell proliferation and cell death upon depletion of BMP signalling. We observed a significant increase in EdU labelled cells in the HFs of the test animals compared to the controls starting from the fifth day till the ninth day, with the peak on the seventh day post tamoxifen injection (compare Figures 1E and 1I with 1F and 1J, respectively and the graph in Figure 1Q). Within the HF, we observed EdU positive cells in the bulge, at the base of HF as well as along the shaft (arrowheads in Figure 1F). On the other hand, we did not observe any cell death, as examined by TUNEL labeling, in the HFs of the test animals till the seventh day after initiating the tamoxifen injection regimen. On the ninth day, however, we observed a significant number of TUNEL positive cells in the HFs of the test animals (compare Figure 1M and 1N, also the graph in Figure 1R). It should be noted that in control animals, no TUNEL positive cells were detected during the entire time window of our investigation.

**Figure 1.**
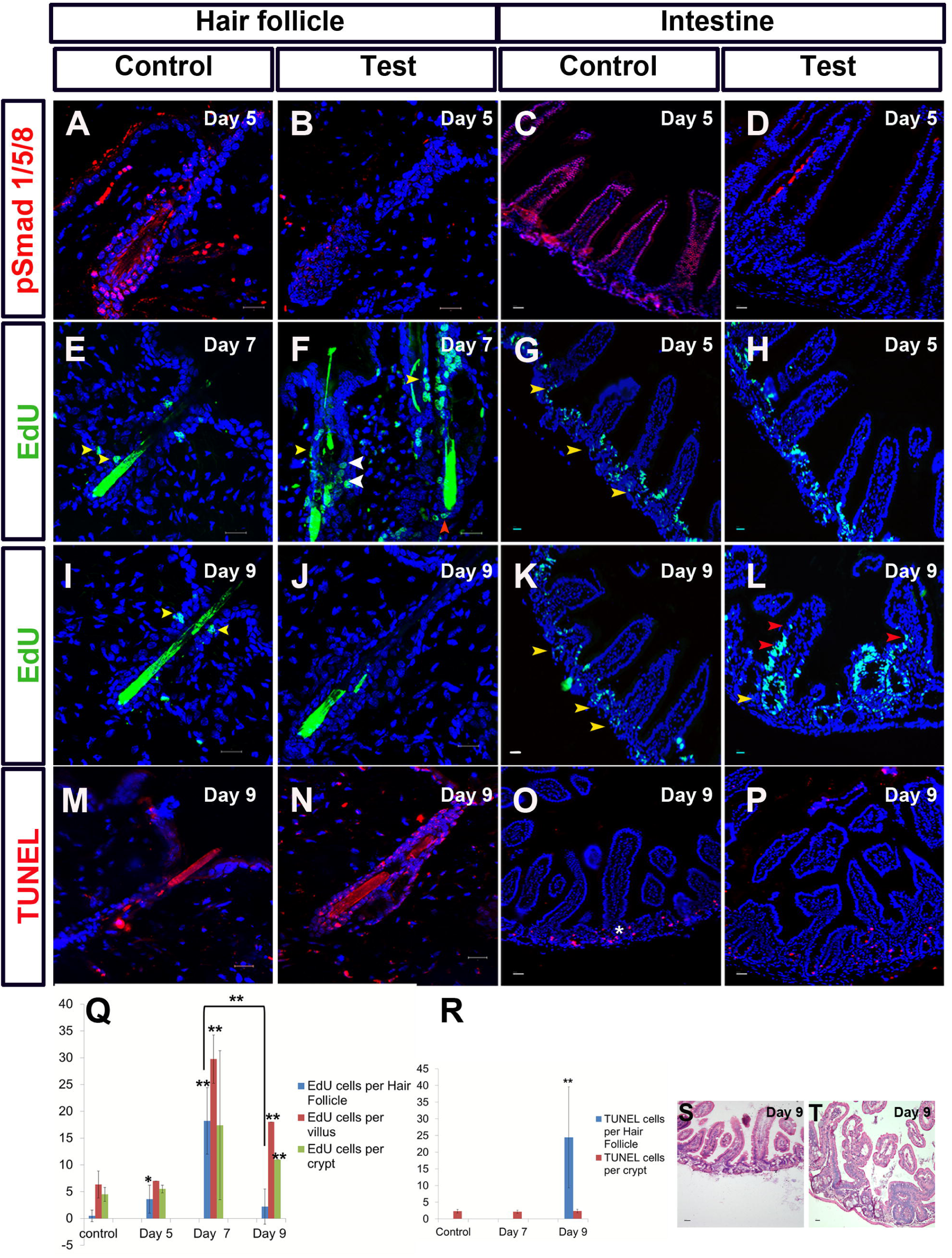
Cellular and histological changes in test animals versus control animals upon depletion of BMP signalling. (A-D) pSMAD 1/5/8 immunoreactivity in the HFs (A-B) and the intestine (C-D). (E-L) State of proliferation in HFs and intestine. (E, F, I and J), EdU incorporation in HFs, EdU positive cells in shafts - yellow arrowheads, in bulge – white arrowheads and in bulb – red arrowheads. (G, H, K and L), EdU incorporation in intestine, EdU positive cells in crypts-yellow arrow heads and in villi-red arrow heads. (M-P) Apoptotic cells in HFs and intestine. (M-N), TUNEL positive cells in HFs; (O-P) TUNEL positive cells in intestine, asterisk shows TUNEL positive cells in crypts of control. Time point indicated on top right corner of each panel refers to the number of days since initiation of tamoxifen injection. (Q) Change in proliferation and (R) change in cell death in HF and intestine following depletion of BMP signalling, * indicates p value< 0.05 and ** indicates p value < 0.005. (S-T) Histology of intestine. Scale bar: 20 μm.

Similar to what we observed in the HFs of the test animals, we observed an increase in EdU positive proliferating cells by the fifth day in the intestinal epithelium as well (compare, Figure 1G and 1H). The maximum increase in proliferation in the intestine was observed by the seventh day (Figure 1Q). However, in contrast to what we observed in the HFs, the number of proliferative cells in the intestine of the test animals did not decrease significantly by the ninth day (compare, Figure 1K and 1L with 1I and 1J, also the graph in Figure 1Q). It is interesting to note that, unlike in control animals where the proliferating cells are restricted to the crypts (arrowheads in Figure 1G and 1K) we could detect many proliferating cells within the villi of test animals (arrowheads in Figure 1L). We would like to highlight the fact that, unlike in the HFs where there is little to no cell death in the resting state, there is normally a basal level of cell death within the intestinal crypts (asterisk in Figure 1O) and this does not increase significantly upon depletion of BMP signalling (compare, Figure 1O with Figure 1P, also the graph in Figure 1R). We also observed distortion in the architecture of the intestinal villi in the test animals by the ninth day where many ectopic crypts were observed (compare Figure 1S and 1T). It is pertinent to mention here that the changes in proliferation and apoptosis closely followed the depletion of BMP signalling. The level of pSMAD 1/5/8went below the detection limit five days after initiating tamoxifen injection followed by the changes in cell proliferation and apoptosis by the seventh day and the ninth day, respectively.

### 2.3. Proliferation as well as survival of hair follicles require a basal level of BMP signalling

It has been previously reported that while BMP signalling is low at the time of anagen induction in HFs, it increases steadily during later stages of anagen and is coincident with the onset of differentiation into various cell types of the HF (Plikus et al., 2008; Schneider et al., 2009). In the set of experiments described above, we observed proliferative cells in the bulge, along the shaft and at the base of HFs suggesting that the depletion of BMP signalling led to induction of anagen. To investigate the relationship between the levels of BMP signalling and proliferation we designed a separate set of experiments this time involving wild type (WT) animals belonging to CByB6F1/J strain. Further, since the changes in proliferation and cell survival were observed in the follicles following global depletion of BMP ligands, to rule out the possibility of non-specific effects, we wanted to use an independent method to locally inhibit BMP signalling in the hair follicle.

It has been shown earlier that repeated shaving causes HFs to go through multiple rounds of cycling (Endou et al.; Hattori and Ogawa, 1983). We therefore first investigated if shaving induces proliferation in the HFs which is a hallmark of anagen. WT males were shaved on the dorsal side to make large shaved patches and the skin was harvested at various time points post shaving. These samples were analyzed for the status of proliferation by assessing EdU incorporation (materials and methods) as well as the BMP signalling by pSMAD 1/5/8 immunoreactivity. Proliferative cells were first observed 48 hours after shaving and we could detect proliferative cells for the next three days which continued upto nine days post-shaving (Figure S2A-S2F and Figure S3). We also observed that the expression of *Shh* and *Lef1*, which are reported to be associated with anagen onset (Kishimoto et al., 2000; Sato et al., 1999), was upregulated within ten days in the HFs from shaved patches as compared to the HFs of the non-shaved skin of the same animals (Figure S4). It was interesting to note that BMP signalling became undetectable by 48 hours after shaving but could be detected again on the fifth day post-shaving (Figure S2G-S2L). This set of experiments demonstrates that induction of proliferation by shaving in resting HFs is also associated with a temporary reduction in the level of BMP signalling. However, this proliferation continues and is observed even when BMP signalling is activated again. In fact we have found that both pSMAD 1/5/8 immunoreactivity and proliferative cells could be detected in the HF up to the ninth day post-shaving (Figure S3).

When we depleted BMP signalling in the test animals by the injection of tamoxifen, we first observed a spurt in cell proliferation in the HFs followed closely by a dramatic increase in cell death (Figure 1). We wondered if this could be explained by the possibility that the newly proliferated HF cells require BMP signalling for their survival. It should be noted that three days after the initial dip that coincides with the onset of proliferation there is a resurgence of BMP signalling after five days of shaving (Figure S2). If our hypothesis is correct, then if we were to deplete BMP signalling in already proliferating follicles, this should advance the onset of apoptosis to a time point earlier than what was observed following Bmp depletion in resting follicles. To test our hypothesis we decided to induce proliferation by shaving. Then, when the cells are already proliferating and BMP signalling is starting to make a comeback (on the fifth day post-shaving) we injected tamoxifen to deplete Bmp2 and Bmp4 proteins (Figure S2L).

We harvested skin samples on the fifth, seventh and ninth day post-shaving (i.e. zeroth, second and fourth day post initiation of tamoxifen injection) (Figure 2A). We did not observe any apoptosis in the samples harvested at the first two time points while we observed many TUNEL positive cells in the sample that was harvested on the ninth day post-shaving (i.e. fourth day post initiation of tamoxifen injection) (Figure 2B). To confirm whether this apoptosis is a result of depletion of BMP signalling, we supplemented the level of Bmp proteins within the shaved patch by subdermal injection of recombinant human Bmp2 (rhBmp2) protein on the eighth day post shaving. We observed that cell death was rescued in the shaved patches supplemented with Bmp protein (Figure 2E). We confirmed that the supplemented rhBmp2 was active by examining pSMAD 1/5/8 immunoreactivity in the samples (compare Figure 2C with 2F, the arrowheads in Figure 2F point to the pSMAD 1/5/8 immunoreactive cells). It should be noted that if we shave the skin and do not deplete BMP activity we do not observe any apoptotic cells in the HFs within the shaved patch but we do detect several cells that are proliferative and pSMAD 1/5/8 immunoreactive (Figure S3). In contrast, in the tamoxifen injected test animals, we could not detect any proliferative cell in the HFs within the shaved patch on the ninth day post-shaving (Figure 2D). Interestingly, we observed that supplementation with rhBmp2 protein led to reappearance of proliferative cells in the HFs on the ninth day post-shaving (compare Figure 2D with 2G). Thus supplementation by rhBmp2 rescues cell proliferation as well as protects against cell death within the Bmp-depleted shaved patches of skin.

**Figure 2.**
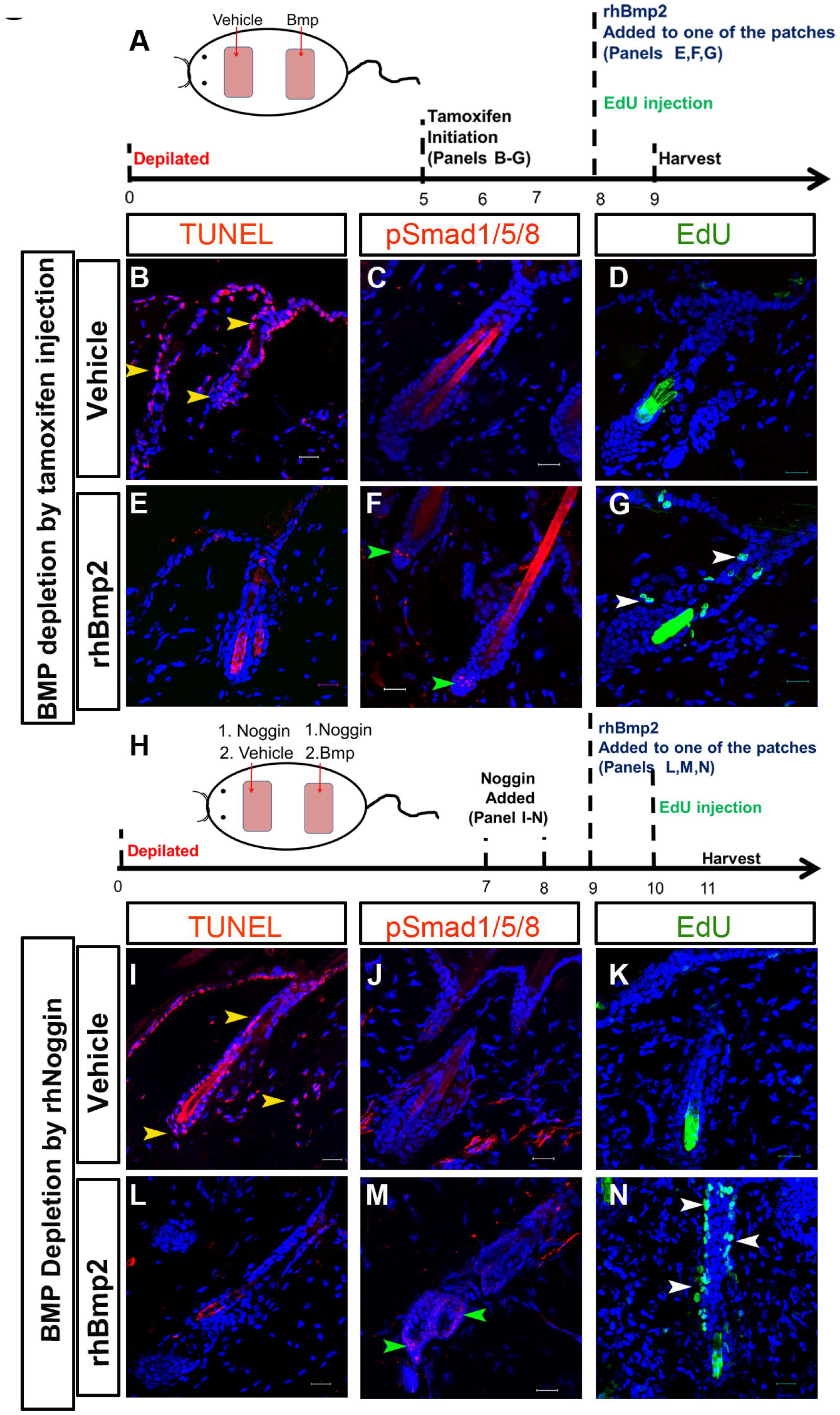
A basal level of BMP signalling is required for proliferation as well as survival of hair follicle cells. (A) Schematic of experimental design. (B-G) BMP signalling depleted by tamoxifen injection in shaved test animals. (B-D) vehicle injected patch and (E-G) rhBmp2 injected patch. (B and E) TUNEL positive apoptotic cells (yellow arrowheads); (C and F) pSMAD 1/5/8 immunoreactivity (green arrowheads) and (D and G) EdU incorporated proliferating cells (white arrowheads). (H) Schematic of experimental design. (I-N) BMP signalling depleted by rhNoggin protein slurry injection in shaved wild type animals. (I-K) Patch with rhNoggin injection followed by vehicle and (L-N) Patch with rhNoggin injection followed by rhBmp2. (I and L) TUNEL positive apoptotic cells (yellow arrowheads); (J and M) pSMAD 1/5/8 immunoreactivity (green arrowheads) and (K and M) EdU incorporated proliferating cells (white arrowheads). Scale bar: 20 μm.

The above set of experiments demonstrates that the depletion of BMP signalling during the proliferative phase advanced the onset of apoptosis by four days. To independently confirm that the cell death observed within the HF of the shaved patch of skin of the test animal is indeed due to depletion of BMP signalling in the HFs and not due to genetic background of the animal or due to the global loss of BMP signalling we devised the following set of experiments.

Wild type males (CByB6F1/J) were used in this series and two patches were shaved in each animal. We subdermally injected recombinant human Noggin protein (rhNoggin) in both the patches on the seventh and eighth day post-shaving. In the first patch (Figure 2I-2K) we injected vehicle only on the ninth and tenth days while the second patch received rhBmp2 protein on the ninth and tenth days (Figure 2L-2N). The samples were harvested 12 hours following the final injection. We observed that depletion of BMP signalling by administration of rhNoggin induced cell death which could be rescued by administration of rhBmp2 (compare Figure 2I with 2L). Indeed, examination of pSMAD 1/5/8 immunoreactivity was consistent with depletion of BMP signalling by rhNoggin which was rescued by rhBmp2 (compare Figure 2J with 2M, the arrowheads in Figure 2M point to the pSMAD 1/5/8 immunoreactive cells). In keeping with the experiment above, proliferation induced by shaving was suppressed in the presence of rhNoggin which could be restored by supplementation with rhBmp2 (compare Figure 2K with 2N).

### 2.4. In BMP signalling depleted hair follicles as well as the intestine presence of active Wnt signalling is not correlated with proliferation

In many contexts of tissue differentiation Wnt signalling has been correlated with cell proliferation (Hauck et al., 2005; Lien and Fuchs, 2014; Tanaka et al., 2011)while active BMP signalling is correlated with cell differentiation (Li and Chen, 2013). In many of these contexts the activation of Wnt signalling often occurs in the absence of BMP signalling (Faigle and Song, 2013; Krausova and Korinek, 2014; Lim and Nusse, 2013; Ray et al., 2015). We observed that nine days after tamoxifen injection, when BMP signalling was depleted in the HFs of the test animals, Wnt signalling was indeed activated (as judged by nuclear β-catenin immunoreactivity) while no activation of Wnt signalling could be seen in the HFs of the control animals (compare Figure 3B to Figure 3A). Similarly, in the shaved skin patches of wild type animals that were subdermally injected with rhNoggin on seventh and eighth days post-shaving, we observed activation of Wnt signalling in the HFs while no such activation could be seen in the shaved patches injected with the vehicle alone (compare Figure 3D to Figure 3C).

**Figure 3.**
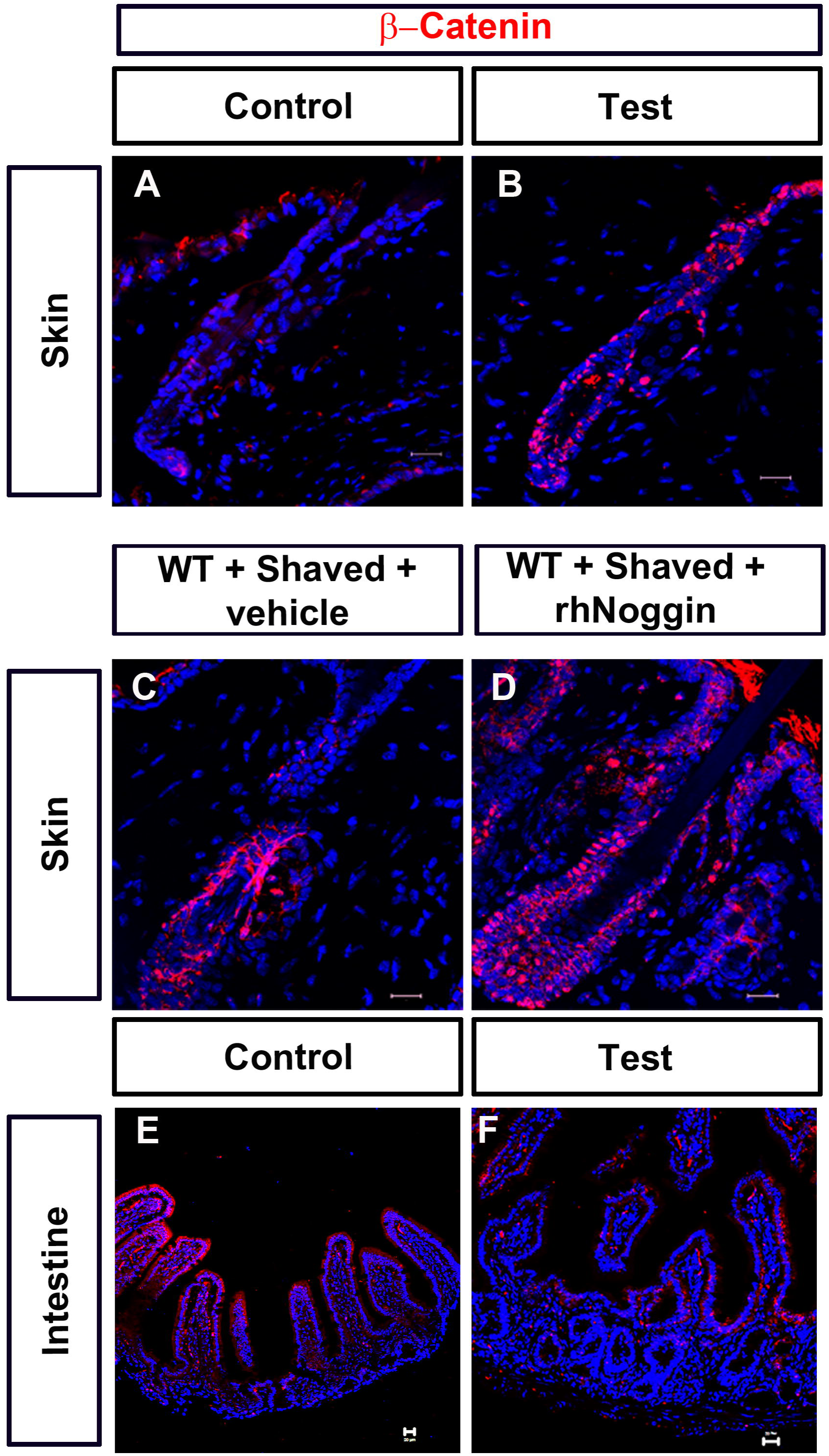
Status of Wnt signalling following BMP depletion in hair follicles and intestine. (A-F) Beta-catenin immunoreactivity. (A) Control and (B) test HFs. (C) HFs in shaved wild type vehicle injected skin patch and (D) HFs in shaved wild type rhNoggin injected skin patch. (E) Control and (F) Test intestine. Scale bar: 20 μm

Previously it has been reported that, during the early stages of anagen, the presence of proliferating cells is correlated with the presence of Wnt signalling in the HFs (Schneider et al., 2009). However, in the test animals we observed that on the ninth day post tamoxifen injection there were no proliferating cells (Figure 1J) even though Wnt signalling was active (Figure 3B). Similarly in the shaved skin patch of wild type animals injected with rhNoggin there were no proliferating cells (Figure 2K) although active Wnt signalling was present in HFs of these patches (Figure 3D).

On the other hand in the context of the intestine we observed a significant increase in proliferating cells both in the crypts as well as the villi of test animals as compared to the control animals (Figure 1), while we could not detect any commensurate increase in Wnt signalling in these structures (compare Figure 3F to Figure 3E).

### 2.5. Depletion of BMP signalling differentially affects the expression of certain differentiation markers in hair follicles

We observed that depletion of BMP signalling initially led to an increase in cell proliferation, which is closely followed by the onset of cell death (Plikus et al., 2008; Schneider et al., 2009). Recently, Genander et. al. demonstrated that BMP signalling is necessary for differentiation of multiple cell lineages of the HF (Genander et al., 2014). Therefore, one explanation for the observed onset of cell death could be that while HFs depleted of BMP signalling are able to proliferate, subsequent lack of BMP signalling, which is required for differentiation, leads them to undergo apoptosis.

To identify if any such differentiation markers may be sensitive to depletion of BMP signalling level, we investigated the expression of a set of some well-characterized markers following manipulation of BMP signalling in the HF. We examined the expression of (i) *Pdgfa* which is a pan-anagen marker and has been suggested to play a role in the maintenance of anagen (Tomita et al., 2006) (ii) *Msx2*, predicted to be a downstream effector of BMP signalling, which is required for keratinocyte differentiation (Botchkarev and Paus, 2003; Jiang et al., 1999; Ma et al., 2003); (iii) Sox9 and K15, outer root sheath (ORS) markers and (v) GATA3, an inner root sheath (IRS) marker.

The expressions of these markers were analyzed in the skin samples of wild type mice where proliferation was induced by shaving. The level of BMP signalling in these shaved patches were manipulated by (i) injecting slurry containing rhNoggin or (ii) injecting rhBmp2 following treatment with rhNoggin for two days. As control we also examined the expression of these markers in shaved patches of skin injected with the vehicle alone. We observed a significant reduction in the expression of *Pdgfa* and *Msx2* in the HFs of rhNoggin injected patches as compared to control patches (compare Figure 4B to 4A and Figure 4F to 4E) which could be rescued by rhBmp2 supplementation (compare Figure 4C to 4A and Figure 4G to 4E). We observed a similar trend with the ORS markers such as K15 and Sox9 (compare Figure 4J to 4I and 4N to 4M; also compare Figure 4K to 4I and 4O to 4M). In contrast, with the IRS marker GATA3, we observed an increase in its expression in the rhNoggin injected patches (compare Figure 4R to 4Q) which could be brought down to the normal level upon supplementation with rhBmp2 (compare Figure 4R to 4Q). To address the concern that some of the observed effects of rhBmp2 supplementation following rhNoggin injection, could be due to independent effects of rhBmp2 we examined the status of expression of these markers in the shaved patches where rhBmp2 was injected alone. We observed that the expression of these markers in HFs of the patches injected with rhBmp2 alone were very similar to that in the HFs in control patches (figure 4D, 4H, 4L, 4P and 4T respectively).

**Figure 4.**
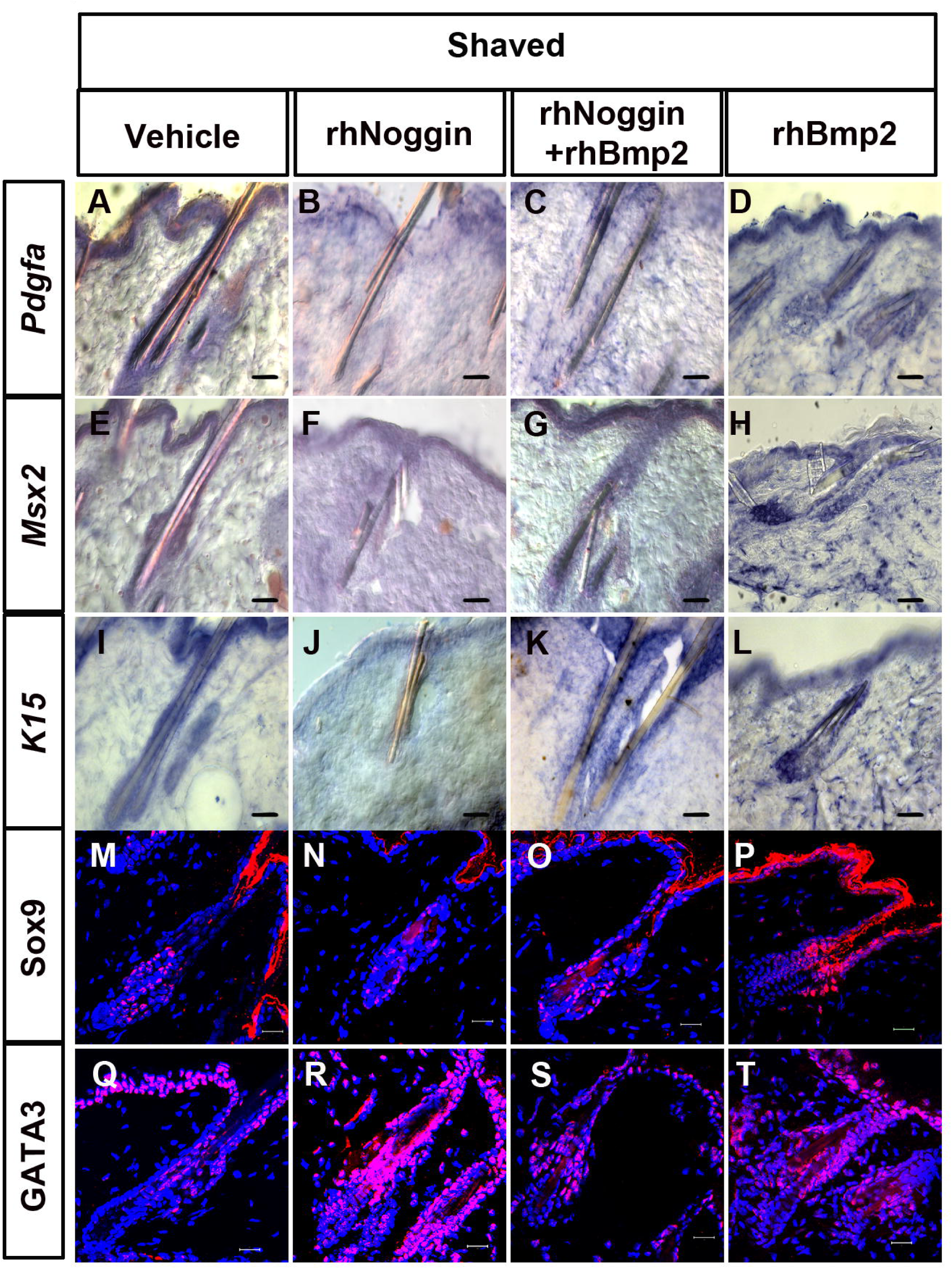
Effect of depletion of BMP signalling on hair follicle differentiation. (A, E, I, M and Q) HFs in shaved patches injected with vehicle; (B, F, J, N and R) HFs in shaved patches injected with rhNoggin; (C, G, K, O and S) HFs in shaved patches injected with rhNoggin followed by rhBmp2 and (D, H, L, P and T) HFs in shaved patches injected with rhBmp2. (A-D), expression of *Pdgfa* mRNA; (E-H) expression of *Msx2* mRNA; (I-L) expression of *K15* mRNA; (M-P) expression of Sox9 protein and (Q-T) expression of GATA3 protein. Scale bar: 20 μm

## 2.6. Differential effects of depletion of BMP signalling in different intestinal cell types

Several histological changes such as the increase in size of the crypts, formation of ectopic crypts, occasional shortening of villi (compare Figure 1T with 1S) and the presence of ectopically proliferating cells upon depletion of BMP signalling (compare Figure 1L with 1K, arrowheads) prompted us to investigate the status of differentiation in the intestine. Two types of epithelial cell lineages emerge from the stem cell niche of the small intestine; secretory and absorptive. Paneth cells, goblet cells and tuft cells are secretory in nature while enterocytes, the major cell type present in the villi, are absorptive in nature. To characterize the changes in the differentiation of these four cell types of the small intestine upon BMP signalling depletion, we investigated the expression of different markers specific for each of these cell types. For detecting goblet cells we used alcian blue staining while for the other cell types we used immunohistochemistry for Sox9 (Paneth cell marker), Doublecortin like kinase 1 (DCLK1, marker for tuft cells) and Villin (enterocyte marker).

We observed that in the crypts of the test animals as compared to the controls there was an increase in the number of the goblet cells (compare Figures 5E with 5A) as well as the Paneth cells (compare Figures 5F with 5B). We also noted an increase in the size of the goblet cells in the test animals (compare Figures 5E with 5A, arrowheads). In contrast, in the test animals we observed a complete absence of DCLK1 positive tuft cells (compare Figure 5G with 5C) as well as significant decrease in the number of Villin expressing enterocytes (compare Figure 5H with 5D). While the number of Villin expressing cells as well as the apparent intensity of Villin immunoreactivity is decreased it should be noted that there was a significant increase in the size of these cells (compare the area enclosed by dotted black lines in Figure 5H’ with 5D’).

**Figure 5.**
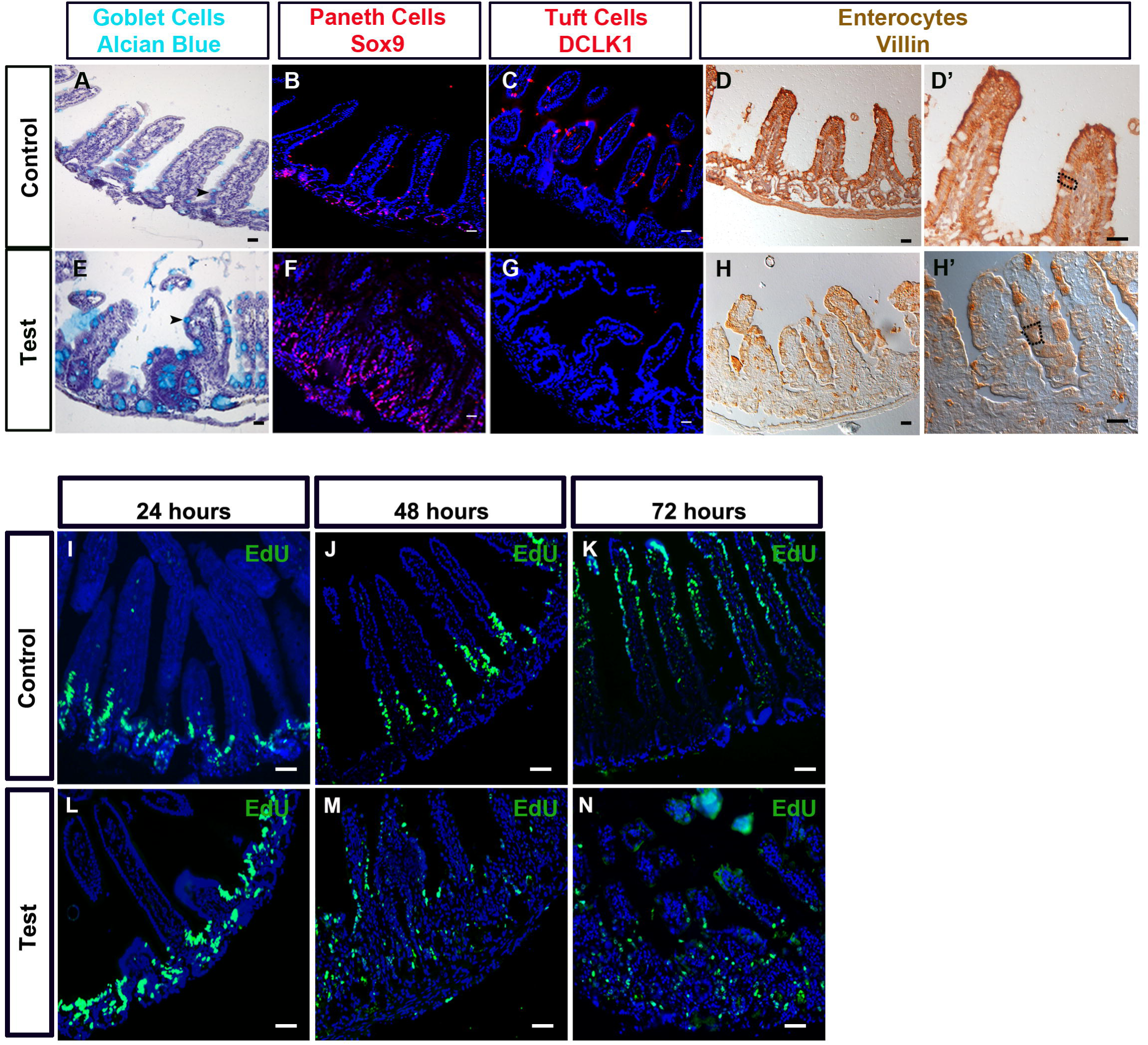
Effect of depletion of BMP signalling on differentiation and migration of intestinal cells. (A-H’) Analysis of intestinal cell type specific markers. (A-D’) Intestine of control animals and (E-H’) intestine of test animals. (A and E) Alcian blue positive goblet cells; (B and F) Sox9 immunoreactivity; (C and G) DCLK1 immunoreactivity and (D, D’, H and H’) Villin immunoreactivity. (D’ and H’) Images at higher magnification of D and H, respectively. Dotted black lines in D’ and H’ outline a single enterocyte. (I-N) Chase of EdU labelled cells in intestine of control and test animals. (I-K) Intestine of control animals and (L-N) intestine of test animals. Location of EdU labelled cells in intestinal epithelium after (I and L) 24 hours; (J and M) 48 hours and (K and N) 72 hours of intraperitoneal EdU injection. Scale bar: 20 μm.

In the intestinal villi cells die by apoptosis at the tip and are replaced by newly born cells that migrate to the tip of the villi from the crypts (Barker, 2014; van der Flier and Clevers, 2009).This process takes place within 3-5 days (Barker, 2014). Thus, the gross changes in villi morphology upon depletion of BMP signalling prompted us to investigate the migration of newly born intestinal cells. In control animals, EdU labelling followed by chase of labelled cells revealed that within 48 hours of EdU labelling most of the cells had left the crypt and populated the lower third of the villi. After 72 hours some of the newly born cells nearly reached the tip of the villi and no EdU labelled cell could be detected in the crypt (Figures 5I – 5K). In contrast, in the test animals even after 72 hours of labelling a large number of EdU labelled cells could be still detected in the crypts and only few could be detected near the tip of the villi (Figure 5N). It should be noted that in the test animals 48 hours post-labelling we observed a large number of EdU labelled cells in the crypts along with many in the villi (Figure 5M). This is in keeping with our observation of ectopic proliferating cells within the villi (Figure 1L).

### 2.7. Depletion of BMP signalling promotes expression of early tumor markers in the intestine

Dysregulation of BMP signalling has been implicated in the pathogenesis of juvenile polyposis syndrome (JPS) which is characterized by, among other hallmarks, ectopic crypt formation (Batts et al., 2006; Haramis et al., 2004). Since we had observed ectopic crypt formation upon depletion of *Bmp2* and *Bmp4* we investigated whether tumor markers are detected in the intestine of the test animals upon depletion of BMP signalling. Thus, we investigated the presence of early tumor markers e.g. pERK1/2and Fatty Acid Synthase (FAS) (Lugli et al., 2006; Visca et al., 1999). In many cells of the intestine of the test animals we have observed nuclear localized p-ERK1/2 (Figure 6B) which has been proposed to be a sign of early neoplastic transformation (Parikh et al., 2012). In contrast, in the intestine of control animals (Figure 6A) most cells had cytoplasmic p-ERK1/2 (compare the *insets* of Figure 6D with 6C). We could also detect many FAS positive cells in the villi epithelium of test animals while no such cell could be detected in the control tissue (compare Figure 6D with 6C).

**Figure 6.**
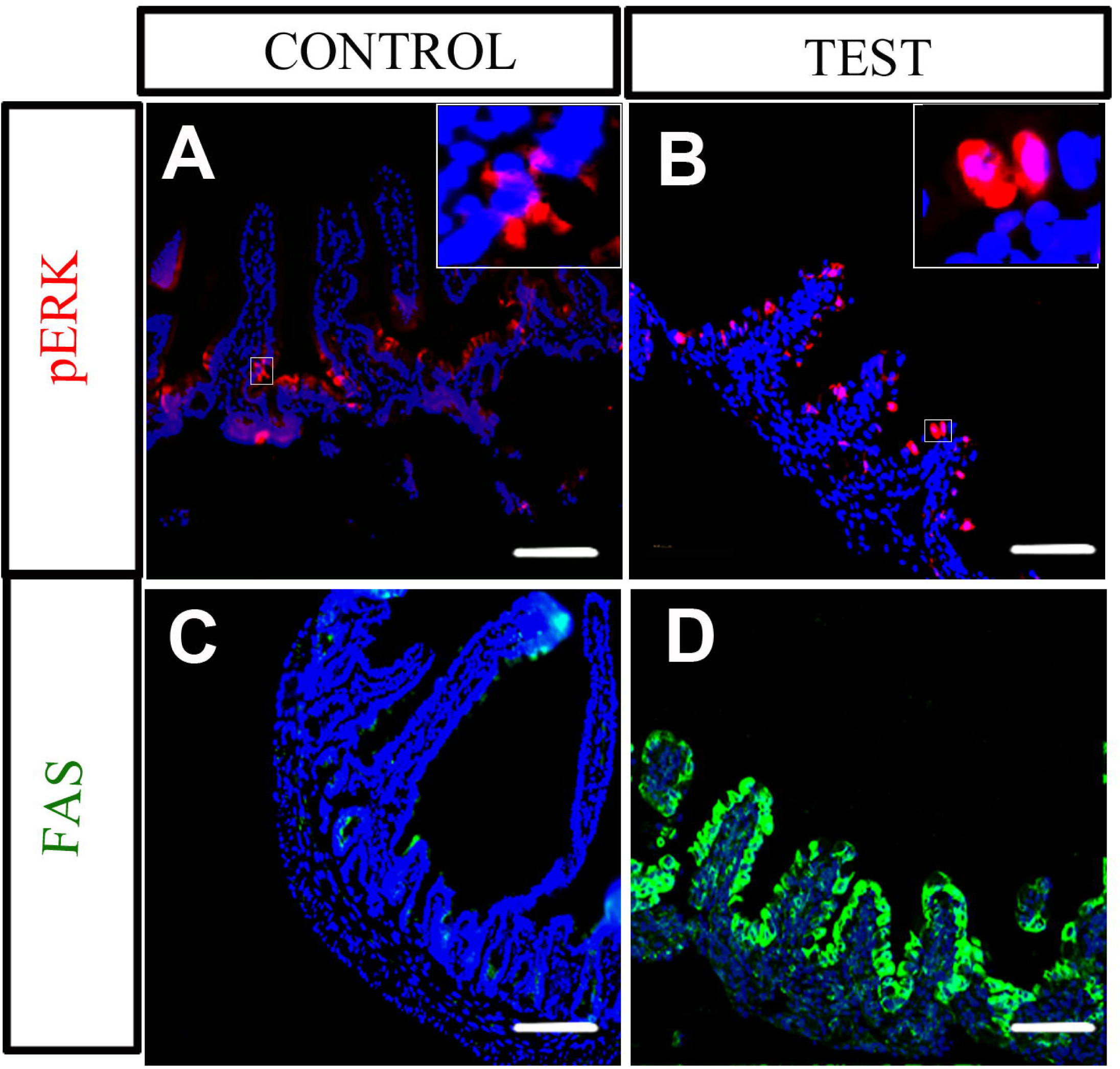
Effect of depletion of BMP signalling on expression of early tumor markers in intestine. (A-B) pERK1/2 immunoreactivity. *Inset*, 2.5 X magnification of the white boxed area. (C-D) FAS immunoreactivity in control and test intestine. Scale bar: 20 μm.

## 3. Discussion

BMP signalling has been studied extensively in the context of embryonic development. However, the role of BMP signalling in adult physiology has not been studied systematically in any model system. Surprisingly, we observed rather acute cellular changes upon ubiquitous depletion of BMP signalling in adult animals. In fact we would like to emphasize that within two days of depletion of BMP signalling in adult animals, homeostasis is significantly perturbed. Such a rapid effect of depletion BMP signalling in adult mice had not been anticipated. In this study, we have focussed on two tissues namely the HFs and the intestine where we observed such rapid changes. It should be noted that embryonic development of these two tissues depends on BMP signalling and both these tissues are known to have relatively high turnover rates (Barker, 2014; Fuchs, 2009; Lee and Tumbar, 2012; Torihashi et al., 2009). Our study demonstrates that homeostasis of these two tissues in adult animals critically depends on BMP signalling. This supports the hypothesis that the same signalling pathway(s) that regulate embryonic development continue to regulate adult tissue homeostasis and this is particularly apparent in tissues with high turnover rates. We have shown that in the HFs, BMP signalling is required for maintaining the HF Stem Cells (HFSCs) in a quiescent state and also for survival and maintenance of proliferative state in HFs. Further, a basal level of BMP signalling is required for maintaining an optimal proliferation rate and differentiation of certain secretory cell types in the intestinal mucosa. Interestingly we found that in the intestine BMP signalling does not seem to be required for cell survival.

Through this study we have demonstrated for the first time that besides depilation, shaving of the dorsal skin of mice is sufficient for inducing certain changes consistent with anagen induction. This was indicated by the initiation of proliferation in hair follicles as well as by the increase in expression of *Shh* and *Lef1*, both of which are known to occur during anagen induction.. Since we observed hair loss following depletion of BMP signalling we investigated the role of this signalling pathway in survival of HF cells. We have conducted several lines of experiments to demonstrate that a resurgence of BMP signalling after the induction of proliferation is critical for the survival of HF cells. It is interesting to note that following depletion of BMP signalling, either by tamoxifen administration or by rhNoggin injection, we observed massive cell death everywhere within the HF including the distal parts where no BMP signalling could be detected even in control HFs (Figure 1). We think that the observed cell death throughout the HF even beyond the normal domain of active BMP signalling may be caused by either of the following reasons. 1) Survival of HF cells that are beyond the domain of active BMP signalling also critically depend on certain diffusible factor(s) whose production is BMP signalling dependent. 2) Our data suggests that proliferative zones within the HF are present in the bottom half which coincides with the domain of active BMP signalling. It is possible that these cells need to initially experience BMP signalling in order to survive even when they have migrated to the distal part of the HF. Thus when BMP signalling is depleted after the tamoxifen injections in test animals the newly born HF cells are not exposed to the initial BMP signalling and thus cannot survive even when they migrate to the distal parts of the HF.

In contrast, in the intestine, depletion of BMP signalling resulted in no significant change in the basal level of apoptosis. Thus our data suggests that the role of BMP signalling in cell survival is highly context dependent.

Our study also highlights context dependent role of BMP signalling in proliferation. It is quite interesting to note that on one hand BMP signalling must be depleted in quiescent HFs to kick-start proliferation while on the other hand a basal level of BMP signalling seems to be necessary to sustain proliferation in HFs. In the intestine depletion of BMP signalling not only causes an increase in the basal level of proliferation in the crypts but also promotes proliferation in an ectopic location such as in the lower portion of the villi where proliferating cells are normally absent. Many examples in existing literature point to the fact that in many tissue contexts, including the HF and the small intestine, presence of BMP and Wnt signalling are mutually exclusive and presence of proliferating cells are correlated with the presence of active Wnt signalling. However, in context of HFs, our data clearly demonstrates that while inhibition of BMP signalling promotes Wnt signalling, it does not lead to proliferation once the initial induction of proliferation is over. This suggests that activation of Wnt signalling may be necessary but not sufficient for promoting proliferation in the HFs. On the other hand in the intestine on depletion of BMP signalling there is a clear increase in cell proliferation as well as presence of proliferative cells in ectopic locations without an associated change in Wnt signalling.

Previous studies where BMP signalling was depleted at embryonic stages have clearly implicated this pathway as a critical player in HF development. However, the severity of the phenotype obtained in these studies and the fact that differentiation occurs at a later stage of development, precluded any analysis of the role of BMP signalling in differentiation of distinct cell types in the HFs. Our strategy of depleting BMP signalling in adult mice allowed us to investigate the possible role of BMP signalling in differentiation of the HF. Presence of proliferating cells is a hallmark of anagen. Depletion of BMP signalling in post-anagen HF abrogates proliferation as well as Pdgfa expression. Taken together our data suggests that BMP signalling is important for maintenance of anagen. Further, marker analysis in Noggin treated shaved HFs suggests that BMP signalling is necessary for ORS differentiation while IRS differentiation per se does not seem to be dependent on the presence of BMP signalling. Our observation vis-à-vis role of BMP signalling in HF differentiation is in-sync with a recent report (Genander et al., 2014)

In earlier studies where BMP signalling was inhibited in different intestinal cell types only the effect on crypt morphology and cell proliferation could be investigated as differentiation was completely blocked (Batts et al., 2006; Haramis et al., 2004). On the other hand in mice with Villin1::Cre mediated conditional knockout of BmprIa a very mild phenotype was observed with differentiation defects only in the Paneth cells (Auclair et al., 2007). In contrast, we observed a dramatic reduction in Villin positive absorptive cells within 4 days of depletion of BMP signalling. We also observed that the differentiation of the secretory cell types such as Paneth cells and Goblet cells increases in the absence of BMP signalling, whereas the differentiation of Tuft cells is decreased. This discrepancy between the results of the earlier study by Auclair *et al* and our study is most likely due to functional redundancy between BMP receptors 1a and 1b whereas we depleted two major Bmp ligands expressed in the intestine. Thus our study for the first time provided an opportunity to assess the role of BMP signalling in differentiation of distinct intestinal cell types. The success of our approach in providing insight into the role of BMP signalling in differentiation of intestinal cells may be attributed to two factors: 1) We decidedly avoided manipulating BMP signalling at embryonic stages and the consequent complications arising thereof and 2) The high turnover rate of the intestinal cells provided the opportunity to assess the immediate effect of depletion of BMP signalling during de novo differentiation of intestinal cell types.

Upon depletion of BMP signalling in the intestine we have observed several cellular changes. The number of proliferating cells is increased, the migration of these newly proliferated cells seems to be affected and ectopic crypts are induced, while apoptosis seems to be unaffected. As a result of all these changes the morphology of the intestine is grossly disrupted, possibly resulting in rectal prolapse. Further, within four days of depleting BMP signalling we observed nuclear pERK1/2 positive cells and a massive upregulation of FAS, both of which are associated with tumorigenesis. In this context it should be noted that BMP loss of function mutations have been associated with JPS which is characterized by tumorigenesis and the formation of ectopic crypts (Batts et al., 2006; Haramis et al., 2004). Thus the mutant mouse generated in this study can serve as a useful model to study the molecular mechanisms underlying JPS.

In conclusion, our strategy to conditionally deplete BMP signalling ubiquitously in adult mice allowed us a glimpse of the variety of contexts in which BMP signalling plays critical roles in maintenance of adult tissue homeostasis. Our approach has highlighted, for the first time, that BMP signalling is needed continuously in adult animals and in absence of which homeostasis is severely affected within a very short span of time, particularly in tissues with high turnover rate.

## 4. Experimental Procedures

### 1.1. Animals strains

*Bmp2*^*C/C*^; *Bmp4*^*C/C*^; *R26CreER/+* (Yadav et al., 2012) mouse strain has been described previously. CByB6F1/J (stock number 100009 Jax labs) were used as the wild type strain.

All animal experiments were conducted according to the protocol approved by the Institute Animal Ethics Committee (IAEC) of Indian Institute of Technology Kanpur, India, which is registered with the Committee for the Purpose of Supervision and Control of Experiments on Animals (CPCSEA).

### 1.2. Tamoxifen inducible conditional depletion of Bmp2 and Bmp4 and genotyping

Six week old *Bmp2*^*C/C*^*; Bmp4*^*C/C*^*; R26CreER/+* mice were administered tamoxifen (Sigma-Aldrich, USA, 2.5 mg/20 g body weight) injection on seven consecutive days. Tissues were harvested at different time points for analysis. The efficiency of recombination was determined using sets of primers described earlier (Prashar et al., 2014). The reduction in Bmp signalling was evaluated by anti-pSMAD 1/5/8 immunoreactivity. All molecular and cellular analyses were conducted with at least 5 animals for each time point. All molecular histological analyses for test and control samples were always performed in parallel on the same slide.

### 1.3. Proliferation Analysis: EdU Labeling

150μg/200μl EdU (Invitrogen C108338) in PBS was administered intraperitoneally 24 hours prior to sacrificing the animals. EdU detection was done with the CLICK-IT EdU imaging kit (Invitrogen C10338).

### 1.4. Cell Death Analysis: TUNEL Detection

Apoptotic cells were detected by TUNEL assay using the cell death detection kit (Roche 12156792910).

### 1.5. Immunohistochemistry

Immunohistochemistry on the skin and intestine samples were done as follows: Skin sections were post-fixed with 4% PFA for 10 minutes, permeabilized with PBT (PBS containing 0.1% Triton X-100, Sigma Aldrich catalog #T8787). Blocking, incubation with primary and secondary antibodies were conducted following standard protocols and counterstained with DAPI. Primary antibodies and dilutions used are as follows: 1:200 anti-β-Catenin (catelog# C7738 Sigma Aldrich), 1:100 anti-Fatty Acid Synthase (catelog# F9554 Sigma Aldrich), 1:300 anti-GATA3 (#HPA 029730 Sigma Aldrich), 1:100 anti-Phospho MAPK (Erk1/2) (#4370 CST), 1:100 anti-pSMAD 1/5/8(#9511 CST), 1:1000 anti-Sox9 (#AB5535, Millipore) and 1:100 anti-Villin (#MAB1671, Millipore).

### 1.6. Depletion of BMP signalling, anagen induction and rescue experiments

The animals were shaved dorsally to make two hairless patches. BMP signalling was depleted using two different methods: (1) Globally in *Bmp2*^*C/C*^; *Bmp4*^*C/C*^; *R26CreER/+* animals by tamoxifen injection starting on the fifth day post-shaving and (2) Locally by sub-dermal injection of 1μg of rhNoggin (Sino Biological Cat#10267-H02H) in CByB6/J animals on the seventh and eighth day post-shaving. BMP signalling level was rescued by injecting 1μg of rhBmp2 (Sino Biological Cat# 10426-HNAE) sub-dermally. A slurry of low melting agarose was used as vehicle for the injection of both proteins (low melting agarose Sigma Cat#A9414). The subsequent injections of proteins were made in the same sub dermal pocket made by the first day injection to ensure that HFs in the same region are targeted every time. The samples of skin were harvested from the shaved patches. The sections were made and only the HFs in direct contact of the subdermal pocket were analysed for the phenotype.

### 1.7. RNA in situ Hybridization in Hair Follicles

RNA in situ Hybridization on 10μm thick frozen skin sections was essentially performed following the protocol described previously (Kawano et al., 2005) with minor modifications.

### 1.8. Imaging

Images were taken using Leica DFC 500 camera attached to Leica DM500B compound microscope. Confocal images were acquired with Zeiss (LSM 780) microscope using DPSS561™ and Chameleon™ lasers. The maximum intensity images were generated using Zen Black™ Software, Carl Zeiss Inc.

### 1.9. Quantification of cell-death and cell proliferation

a. **Hair follicles:** Cell death was estimated by counting the total number of TUNEL positive cells present per HF in 10 randomly selected fields from each skin sample of five independent test and control animals for each time point. Cell proliferation was estimated by counting EdU positive cells in a similar manner.
b. **Intestine:** Cell death was estimated by counting the number of TUNEL positive cells per crypt from five randomly selected fields from each intestine sample of five independent test and control animals. Cell proliferation was estimated by counting the number of EdU positive cells present per crypt or per villi in each field. The mean and standard deviation in each case was calculated using MS Excel. Two tailed T test was used for calculating statistical significance.

## Authors Contributions

A.B. conceived the study; A.B., J.S. and A.N. developed the experimental approach; A.N. and A.L.N. performed the HF analyses; M.K. optimized RNA in situ protocol for HFs and R.G. repeated the protocol; P.N. and A.N. performed the intestinal analyses; A.B., J.S., P.N. and A.N. wrote the manuscript.

## Acknowledgements

We acknowledge Prof. E. Fuchs for providing us with the antibodies for HF markers. This work was supported by a grant (SR/SO/AS-16/2010) from DST. AN and PN were supported by grants from UGC and DBT, respectively. Affiliation of ALN with Abhilash Chemicals has no conflict of interest with the data generated and presented here.

